# Automated patch clamp screening of amiloride and 5-N,N-hexamethyleneamiloride (HMA) analogs identifies 6-iodoamiloride as a potent acid-sensing ion channel inhibitor

**DOI:** 10.1101/2022.03.12.484055

**Authors:** Rocio K. Finol-Urdaneta, Jeffrey R. McArthur, Ashraf Aboelela, Richard S. Bujaroski, Hiwa Majed, Alejandra Rangel, David J. Adams, Marie Ranson, Michael J. Kelso, Benjamin J. Buckley

**Affiliations:** Electrophysiology Facility for Cell Phenotyping and Drug Discovery, Wollongong, NSW 2522, Australia; Illawarra Health and Medical Research Institute, Wollongong, NSW 2522, Australia; School of Chemistry and Molecular Bioscience, University of Wollongong, NSW 2522, Australia; Molecular Horizons, University of Wollongong, NSW 2522, Australia

**Keywords:** Acid-sensing Ion Channel, ASIC, ASIC inhibition, Amiloride, small molecule, Synchropatch

## Abstract

Acid-sensing ion channels (ASICs) are transmembrane sensors of extracellular acidosis and potential drug targets in several disease indications, including neuropathic pain and cancer metastasis. The K^+^-sparing diuretic amiloride is a moderate non-specific inhibitor of ASICs and has been widely used as a probe for elucidating ASIC function. In this work, we screened a library of 6-substituted and 5,6-disubstituted amiloride analogs using a custom-developed automated patch-clamp protocol and identified 6-iodoamiloride as a more potent ASIC1 inhibitor. Follow-up IC_50_ determinations in tsA-201 cells confirmed higher ASIC1 inhibitory potency for 6-iodoamiloride **97** (hASIC1 **97** IC_50_ 88 nM cf. amiloride **11** IC_50_ 1.7 μM). A similar improvement in activity was observed in ASIC3-mediated currents from rat small diameter dorsal root ganglion neurons (rDRG single-concentration **97** IC_50_ 230 nM cf. **11** IC_50_ 2.7 μM). 6-iodoamiloride represents the amiloride analogue of choice for studying the effects of ASIC inhibition on cell physiology.

---

Acid-sensing ion channels (ASICs) are proton-gated members of the ENaC/Degenerin superfamily; a diverse group of trimeric transmembrane ligand-gated ion channels found across the chordate phyla.^1, 2^ Six ASIC isoforms have been identified in mammals. These include splice variants ASIC1a/b and ASIC2a/b and the discretely encoded ASIC3 and ASIC4, all of which are expressed predominately in neuronal tissues.^3^ Isoform subunits associate into homo or heterotrimeric channels that differ in *p*H sensitivity, activation kinetics, ion selectivity and pharmacology depending on subunit composition and stoichiometry.^7^ ASICs function primarily as transmembrane acid sensors, allowing the selective influx of Na^+^ and Ca^2+^ (ASIC1a/b) in response to small changes in extracellular *p*H.^4, 5^ ASICs affect action potential generation and conductance by modulating neuronal membrane depolarization. To date, ASIC function has been implicated in synaptic plasticity and learning, fear and anxiety behaviors and pain sensation.^6^

Despite evidence suggesting roles for ASICs in multiple disease states, relatively few small molecule inhibitors and clinical studies have been reported (**Figure 1**).^7^ An early ASIC program from Abbott identified 2-naphtamidine A-317567 **1**, which showed significant efficacy in rat models of complete Freund’s adjuvant (CFA)-induced chronic inflammatory and post-operative pain.^8^ Merck reported a novel series of 2-arylindoleamidines (e.g. **2**) with higher ASIC potency and efficacy similar to naproxen in the rat inflammatory pain model.^9^ Efforts aimed at improving oral bioavailability by replacing the amidine produced 6-substituted 2-chlorobenzothiophenmethanamines (e.g. **3**), which showed high ASIC3 potency, efficacy similar to naproxen in pain models and good CNS penetrance.^10^ Ultimately, **3** showed poor PK in dogs and off-target promiscuity, preventing its further development.^10^ Merck also reported a series of A-317567 analogs (e.g. **4**) with improved ASIC3 potency and anti-nociceptive efficacy superior to naproxen in osteoarthritis rat models.^11^ Other work from Merck addressed the difficulty of lead discovery for ASICs by developing high-throughput workflows for automated patch-clamp screening, allowing the screening of the Merck Fragment Library at high concentration (up to 2 mM).^12^ Initial library screenings followed by structure-based similarity searching produced 12 distinct hit chemotypes with activities in the 9-1000 μM range. Further optimization of the 4-aminopyridine, 2-aminopyridine (e.g. **5**) and tetrahydroisoquinoline classes produced leads with low μM activity, low LE and good rat PK, although these were not evaluated in rat pain models due to high plasma protein binding. More recently the tetrahydroisoquinoline NS383 **6**, was reported as a potent and selective inhibitor of rat ASIC1a and ASIC3-mediated currents.^13^ NS383 showed efficacy superior to amiloride in rat models of inflammatory and neuropathic pain. Buta *et al*., 2016 reported a highly potent 2-oxo-2H-chromene-3-carboxamidine derivative 5b, **7**, (ASIC1a IC_50_ = 27 nM) effective in decreasing long-term potentiation currents associated with epilepsy in rat CA3-CA1 hippocampus.^14^ 5b was developed with consideration of key interactions between the RRR motif of Psalmotoxin 1 (PcTX1), a potent peptide toxin know to target the extracellular *p*H sensor domain of ASIC^15^, and the orthosteric extracellular domain of the protein. The inhibitory activity of 5b showed a profound dependence on *p*H, with a 3-order of magnitude decrease in potency observed at *p*H 5.0 relative to pH 6.7, suggesting competition between inhibitor and H^+^ within the *p*H sensor domain. Dose-dependent effects on steady-state ASIC1a current activation and decreased maximum evoked responses suggested a negative allosteric modulatory profile of activity. Diminazene **8** the common livestock anti-protozoal drug was the most potent diarylamidine tested, with an IC50 of 280 nM for ASIC-mediated current in mouse primary hippocampal neurons.^16^ A recent phenotypic screening and medicinal chemistry optimization effort starting from diminazene has described a series of dihydropyrrolo[3,4-*d*]pyrimidinedimethylpyrrazole inhibitors selective for ASIC2.^17^ While showing only moderate potency, the lead from this series 2u **9** (ASIC2a IC_50_ = 17.4 μM), featured an excellent PK profile and high CNS penetrance. Of the small molecule ASIC inhibitors described to date, PPC-5650 (**10**, core structure of class inferred from patent literature^18^, specific structure not disclosed), a potent ASIC1a-selective methylquinoline-based inhibitor is the only candidate to have progressed to clinical trials, reducing hyperalgesia readouts in Phase I studies of UVβ-induced sunburn^19^ and gastroesophageal reflux disorder (GERD).^20^

**Figure 1.**
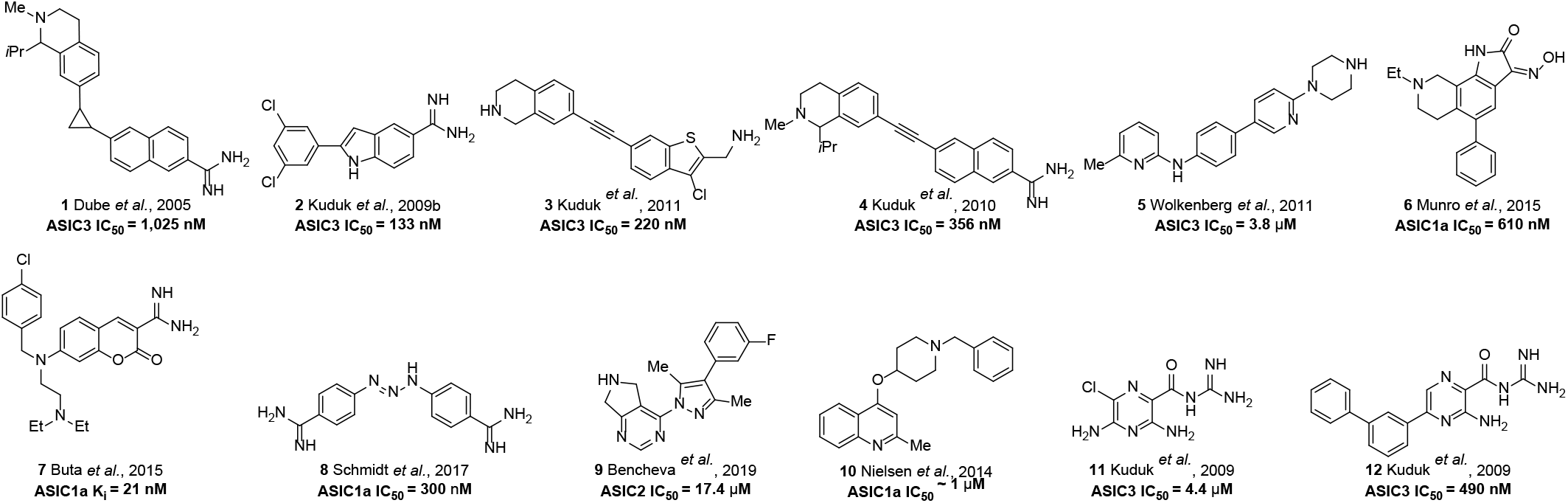
Small molecule inhibitors of ASIC.

Amiloride **11** is an oral K^+^-sparing diuretic used in the management of hypertension and congestive heart failure that exerts its effects through inhibition of renal epithelial sodium channels (ENaC).^21, 22^ Amiloride is a promiscuous drug known to bind to a diverse range of molecular targets, including GPCRs^23–25^, transmembrane ion channels^26–31^ and trypsin-like serine proteases^32, 33^. Despite its lack of selectivity, amiloride has a good safety record with dose-related ENaC-mediated hyperkalemia the only notable risk.^34^ Amiloride is a modest non-selective ASIC inhibitor (IC_50_ = 3-30 μM) that shows anti-nociceptive effects at relatively high doses in rodent models of chronic pain and in acid-induced nociception in humans.^35, 36^ Researchers at Merck produced a library of 5-substituted 6-hydroamiloride analogs (e.g. **12**) that showed up to 8-10-fold improved potency against the peripheral pain target ASIC3 and efficacy equivalent to naproxen in inflammatory pain models.^37^

We recently reported on the activity of two novel series of 6-substituted amiloride^38^and 5-*N*,*N*-hexamethylene amiloride^33^ analogs as inhibitors of uPA; a trypsin-like serine protease linked to cancer metastasis^39^, rheumatoid arthritis^40^ and other inflammatory diseases.^41, 42^ Lead inhibitors from these efforts showed low nM uPA potency, high target selectivity across the serine hydrolase superfamily, loss of unwanted ENaC activity, low hERG activity and *in vivo* anti-metastatic effects in xenografted mouse models of lung and pancreatic cancer.^33, 38, 43^ In this work, we characterise the structure-activity relationships of compounds from these series and related amiloride derivatives as inhibitors of proton-activated currents mediated by ASICs using automated patch clamp electrophysiology, identify an analog with improved potency, and test its potential as pharmacological tool to probe endogenous human and rodent ASIC function.

## Results and Discussion

### Assay development

Acid-sensitive currents mediated by ASIC1 channels have been described in HEK-293 human embryonic kidney cells.^44^ We initially assessed *p*H-sensitive (ASIC) currents in tsA-201 (an SV40 T antigen-transformed HEK-293 clone) for automated patch-clamp (APC) screening of large compound libraries as a low cost alternative to proprietary ASIC-overexpressing cell lines. From a conditioning *p*H (*p*H_C_) 7.4, tsA-201 cells exposed to acidic stimuli (*p*H_S_= 5.5) displayed robust inward whole-cell currents (35.2 ± 3.1 pA/pF, n = 20) that activated monoexponentially with a time constant of 81.2 ± 4.9 ms (n = 20) under APC (holding potential, V_h_ = −60 mV). Currents elicited by 1 s long exposure to increasing proton concentrations from *p*H 8.0 to *p*H 5.0 (*p*H_C_ 7.4 applied between stimuli, **Figure 2A**) were quantified by integration of the area under the current signal reflecting the total charge inwardly displaced by the stimuli and used to determine the *p*H at which 50% of the maximal charge could be elicited or half-activation *p*H (*p*H_0.5_). The *p*H dependence of steady-state desensitization (SSD) was determined by 1 min pre-exposure to a series of *p*_C_ solutions followed by a 1 s application of *p*H 5.5 solution (**Figure 2B**; *p*H_C_ and *p*H_S_ are marked by thick and thin bars, respectively) resulting concentration-response curves rendered an activation *p*H_0.5_ of 5.9 ± 0.04 (n = 6) and a SSD *p*H_0.5_ of 6.5 ± 0.03 (n = 8) (**Figure 2C**). Confocal micrographs from tsA-201 cells stained with a pan-ASIC antibody cocktail revealed discrete foci of immunostaining that overlapped with wheat germ agglutinin counterstaining of the plasma membrane (**Figure S1 E-H**) supporting the expression of ASIC channels in this cell line. Furthermore, proton activated currents elicited by exposure to *p*H 5.5 were inhibited by the ASIC1-selective peptide toxin Mambalgin 1 [Diochot, 2012 #162] with a *K_D_* of 27.7 ± 2.8 nM (n = 6) (**Figure 2D**); and were absent upon siRNA knockdown of ACCN2 transcripts encoding for ASIC1 (GFP^CTR^ = −36.7 ± 3.2 pA/pF, n = 6; GFP^-ACCN2^ = −5.2 ± 3.4 pA/pF, n = 6; unpaired t test p = <0.0001). (**Figure 2E**). Positive immunostaining for the canonical isoform ASIC1a (**Figure S1 A-D**) that correlated with the pattern observed using the pan-ASIC1 MaB cocktail (**Figure S1 E-H**) and the lack ASIC3 signal, are consistent with previous reports^44^ supporting ASIC1a channels as the primary ASIC isoform mediating acid-sensitive currents in HEK-293 cells. Thus, the functional characterization of ASIC mediated currents in tsA-201 cells supports their use as a readily available and low-cost option for library screening at physiologically relevant expression levels of ASIC expression.

**Figure 2.**
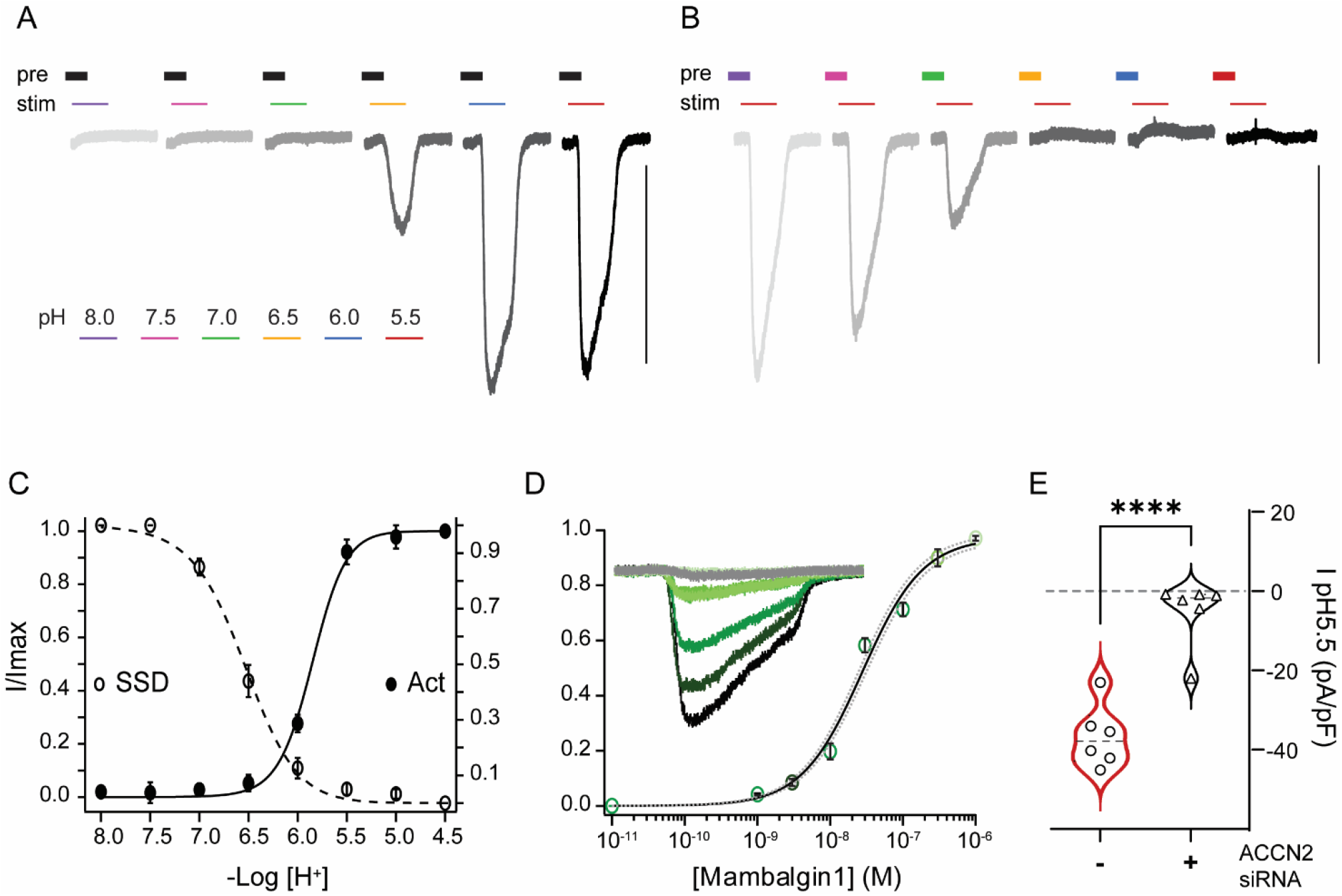
Characterization of endogenous *p*H-sensitive inward currents in tsA-201 cells. **A)***p*H activation dependence, **B)** steady-state *p*H deactivation dependence. The *p*H of the 1 s stimulus application corresponds to colors shown in legend in A. The *p*H of the conditioning solution is indicated by the thick bar above: **A)** *p*H 7.4 (black); **B)** *p*H as per legend in **A)**, **C)**half-maximal activation (*p*H_0.5_ = 5.9 ± 0.04; n = 6) and steady-state deactivation (SSD *p*H_0.5_ = 6.5 ± 0.03l n = 8) and **D)** Inhibition of proton-sensitive inward currents (*p*H 5.5) in the presence of Mambalgin 1 (Mb1, IC_50_ of 27.7 ±2.8 nM; n = 6). The inset shows currents activated by a *p*H jump of 5.5 in the presence of progressively higher [Mb1] (dark to light green). **E)**Quantification of currents elicited by a *p*H 5.5 in tsA-201 cells transfected with GFP (red, I_*p*H5.5_ = −36.7 ±3.2 pA/pF, n = 6) and cells transfected with GFP and siRNA against ACCN2 transcripts (black, I_*p*H5.5_ = −5.2 ±3.4 pA/pF, n = 6; unpaired *t*-test p = <0.0001).

The potency of >90 5- and 6-substituted amiloride analogues against ASIC1-mediated currents in tsA-201 cells was determined by an automated patch clamp protocol using the SyncroPatch 384PE and compared to that of amiloride. The assay design allowed for parallel testing of 9 analogs and amiloride **11** on each 384-well chip (**Figure S3**). When tested at 30 μM, amiloride and HMA inhibited 88 and 79% of ASIC1-mediated of *p*H5.5 elicited current respectively (**Table 1**). Substitution of amiloride and HMA with 6-(hetero)aryl substituents invariably led to decreases in activity (1.1 to 4-fold) relative to the parent compounds (**Tables 1&2**). In general, only modest differences were seen between the amiloride analogs and their matching HMA equivalents bearing the same 6-substituent.

**Table 1.**
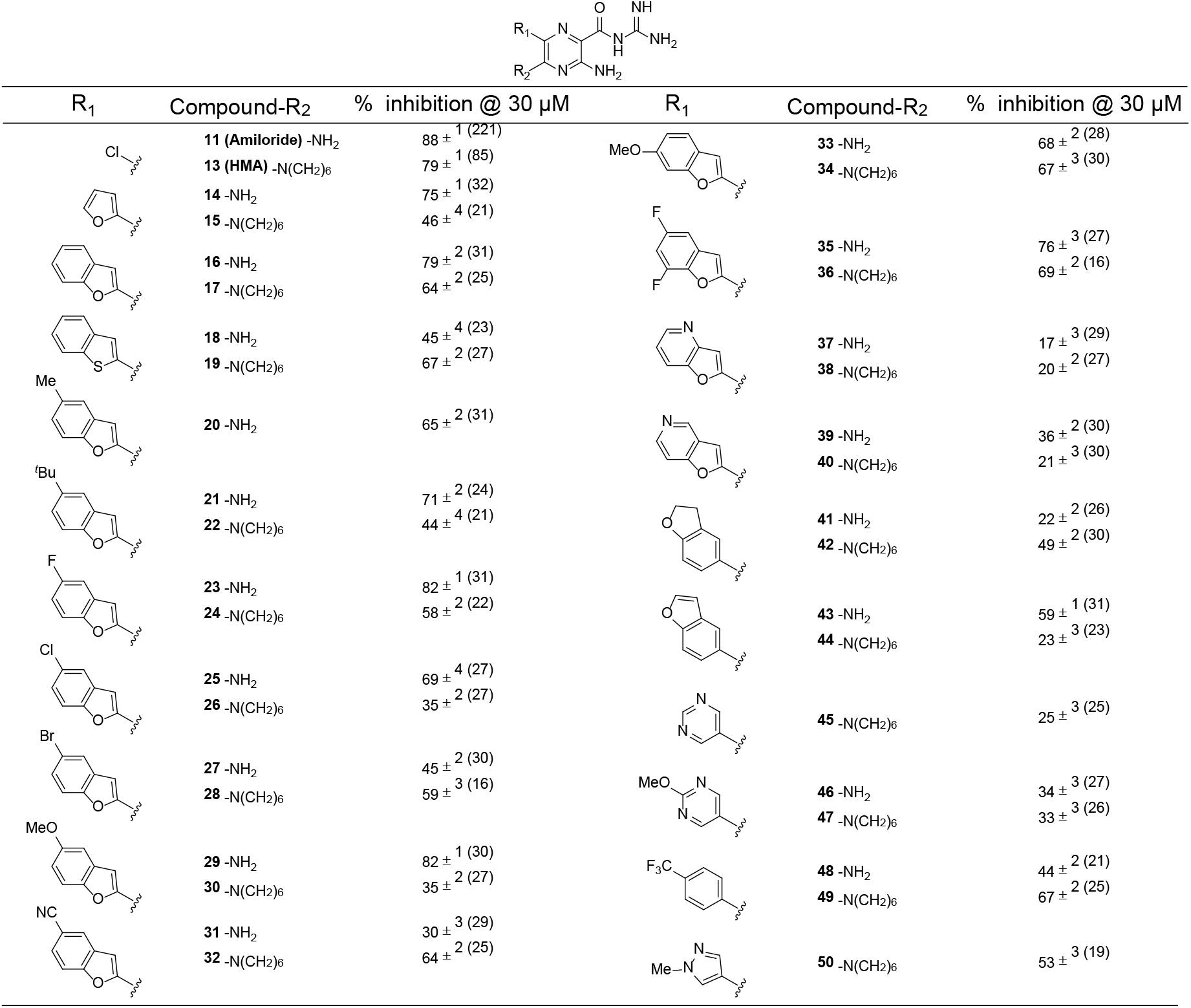
Inhibition of ASIC1 by 6-(hetero)aryl substituted amiloride and HMA analogs. Values represent the mean ± SEM, with number of technical replicates shown in parentheses.

**Table 2.** Additional 6-substituted HMA analogs.

| Compound | R | % inhibition @ 30 μM |
| --- | --- | --- |
| 51 | | 64 ± <sup>2</sup> (30) |
| 52 | | 60 ± <sup>3</sup> (20) |
| 53 | | 48 ± <sup>3</sup> (20) |
| 54 | | 44 ± <sup>4</sup> (19) |
| 55 | | 53 ± <sup>3</sup> (19) |
| 56 | | 70 ± <sup>2</sup> (27) |
| 57 | | 25 ± <sup>3</sup> (16) |
| 58 | | 23 ± <sup>3</sup> (23) |

Previous reports of positive SAR at the 5-position of amiloride motivated screening of analogs containing the benzofuran-2-yl group at position 6 and variously substituted amines at position 5 (**Table 3**).^37^ In all cases, substitution of the amino group reduced potency relative to the parent 6-(benzofuran-2-yl) amiloride analog **16**. Interestingly, reduced activity was observed with groups previously shown to improve the potency of amilorides against ASIC3, (e.g. 5-benzylamine **65** and 5-phenylethylamine **66**).^37^

**Table 3.** Analogs of compound **16** containing substituted amines at position 5.

| Compound | R | % inhibition @ 30 μM |
| --- | --- | --- |
| 59 | | 59 ± 2 (31) |
| 60 | | 54 ± 2 (27) |
| 61 | | 66 ± 2 (25) |
| 62 | | 46 ± 2 (32) |
| 63 | | 22 ± 2 (25) |
| 64 | | 10 ± 3 (30) |
| 65 | | 39 ± 3 (28) |
| 66 | | 31 ± 4 (24) |

The weakly basic acylguanidine of amiloride is a critical determinant of affinity for several ion channel and protease targets.^45^ X-ray co-crystal structures of amiloride bound to uPA^46^ and chicken ASIC1a^47^ showed that the binding contributions from the acylguanidine group stem from interactions with the charged sidechains of acidic glutamate residues. To establish whether the basic acylguanidine group was required for ASIC1 activity in these series, we screened analogs based on **16** featuring other groups at the 2-position (**Table 4**). Methyl ester **67** and the corresponding free acid **68** showed considerably lower activity than **17**, as did the 2-*N*,*N*-dimethylamide **69**. Similar decreases in activity were seen with compounds containing 5-membered heterocyclic acylguanidine isosteres; e.g. 5-amino-1,2,4-triazole **70** and 3-amino-1,2,4-oxadiazole **71**. A smaller loss of activity was observed with the 3-*N*,*N*-dimethyl-1-propanone analog **72**. Together these results suggest that an ionizable basic moiety is required at 2-position of the shared pyrazine core for ASIC inhibition.

**Table 4.**
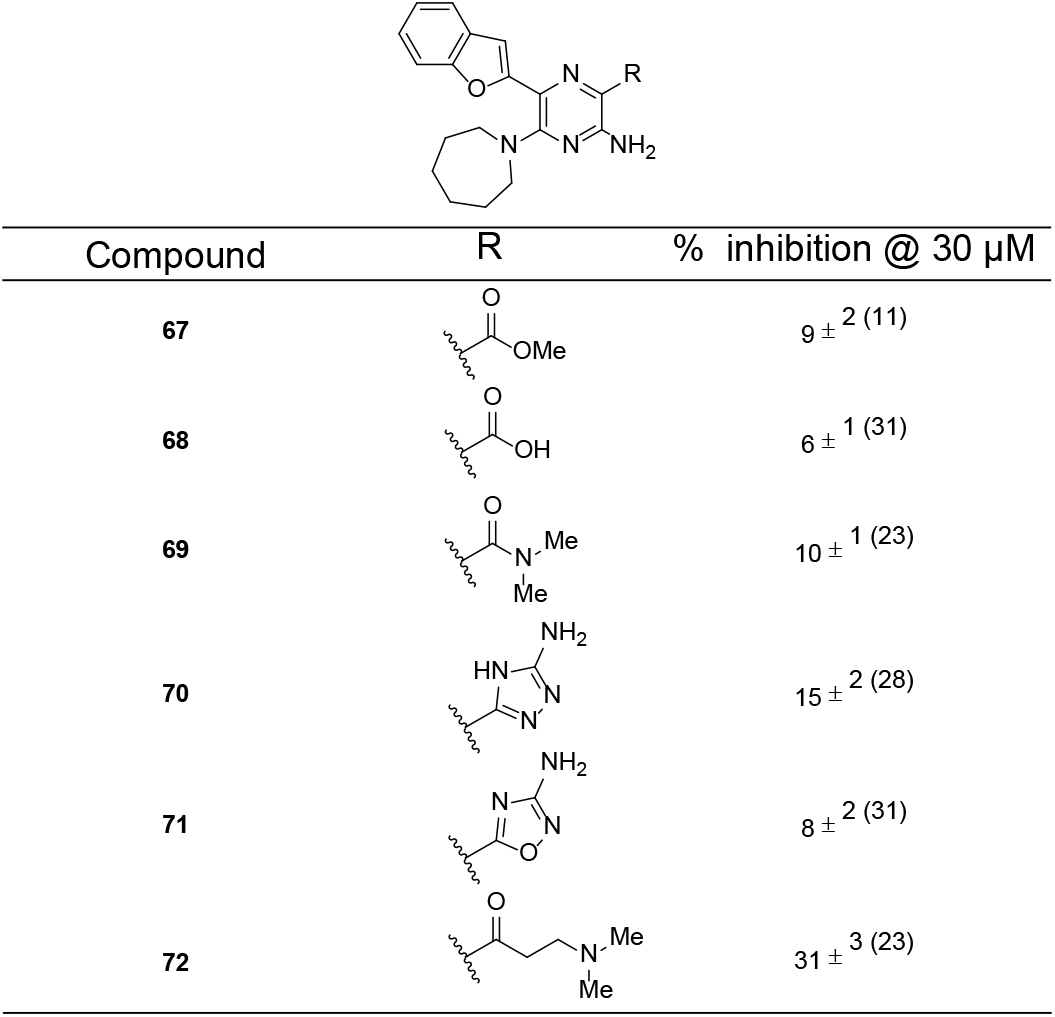
Analogs of **17** containing acylguanidine replacements

| Compound | R | % inhibition @ 30 μM |
| --- | --- | --- |
| **67** | | $9 \pm 2$ (11) |
| **68** | | $6 \pm 1$ (31) |
| **69** | | $10 \pm 1$ (23) |
| **70** | | $15 \pm 2$ (28) |
| **71** | | $8 \pm 2$ (31) |
| **72** | | $31 \pm 3$ (23) |

A series of 4-amino substituted analogs of the 6-pyrimidine-**47** (**Table 5**) showed that replacing the 4-MeO group with various amino groups was not tolerated, decreasing activity by up to 5-fold. Replacing the 5-azepane with the smaller pyrrolidine group while maintaining the 4-methoxypyrimidine at the 6-position **77** (**Table S1**) did not decrease activity relative to **47**.

**Table 5.**
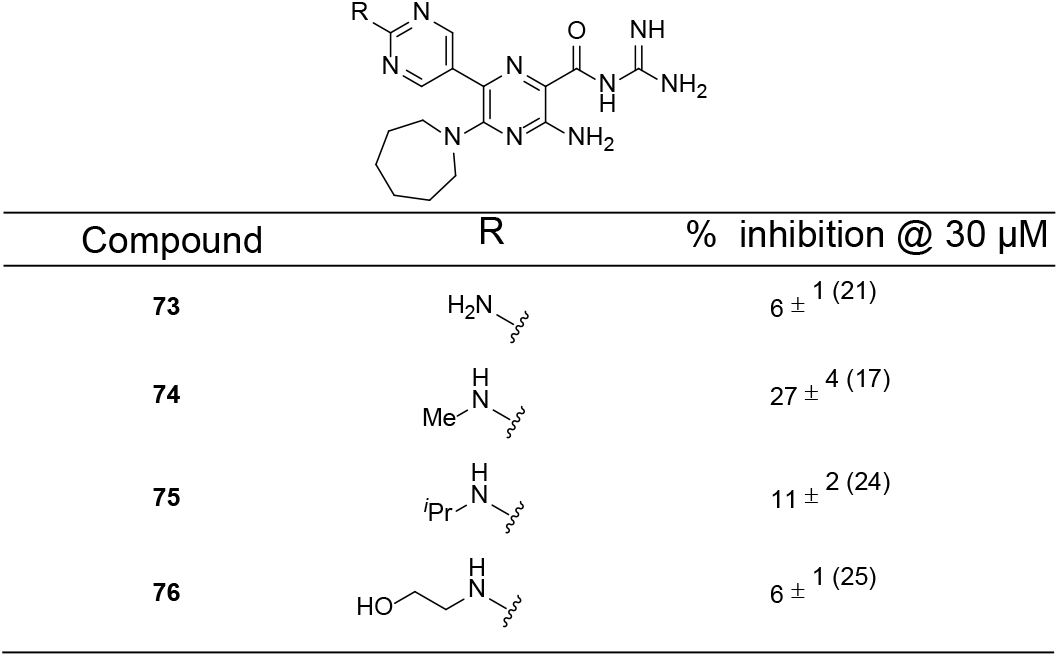
Analogs containing 4-substituted pyrimidines at position 6.

| Compound | R | % inhibition @ 30 μM |
| --- | --- | --- |
| **73** | | $6 \pm 1$ (21) |
| **74** | | $27 \pm 4$ (17) |
| **75** | | $11 \pm 2$ (24) |
| **76** | | $6 \pm 1$ (25) |

Analogues containing the 4-CF3phenyl group at position 6 and pyrrolidine **78** or cyclopropyl **79** groups at position 5 showed slightly improved potency relative to the 5-NH2-substituted counterpart **48** and similar potency to the 5-azepane analog **49** (**Table 6**).^37^ The 5-morpholino amiloride **80** showed slightly lower activity than amiloride, similar to what had been reported earlier for this compound against ASIC3 (**Table 7**).^37^ 6-Substituted 5-morpholino derivatives 6-(pyrimidin-5-yl) **81** and 6-(benzofuran-2-yl) **82**, showed similar activity to their 5-azepane counterparts. Attaching a 1,4-oxazepane ring to position 5 reduced activity in all analogs **83-86** relative to the corresponding 5-azepane equivalents (**Table 8**).

**Table 6.**
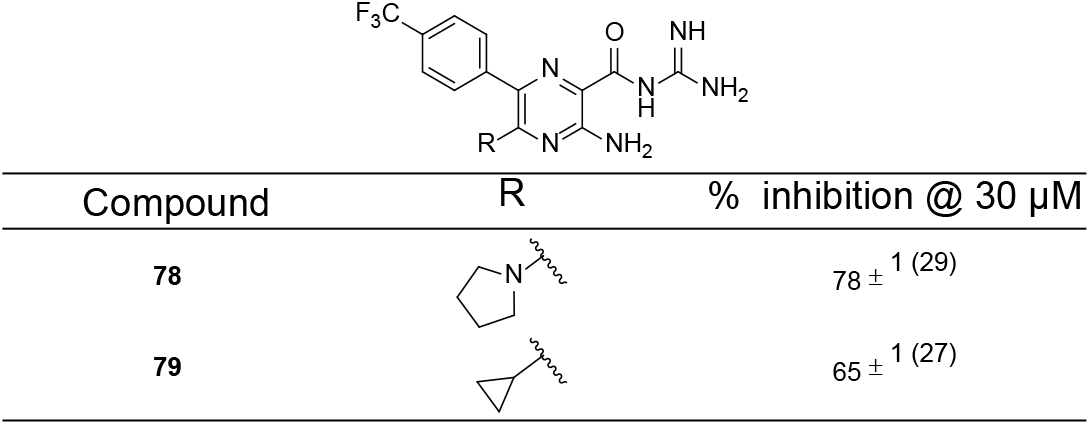
Additional analogs containing 4-F_3_phenyl groups at position 6.

|  |  |  |
| --- | --- | --- |
| Compound | R | % inhibition @ 30 μM |
| **78** | | 78 ± 1 (29) |
| **79** | | 65 ± 1 (27) |

**Table 7.**
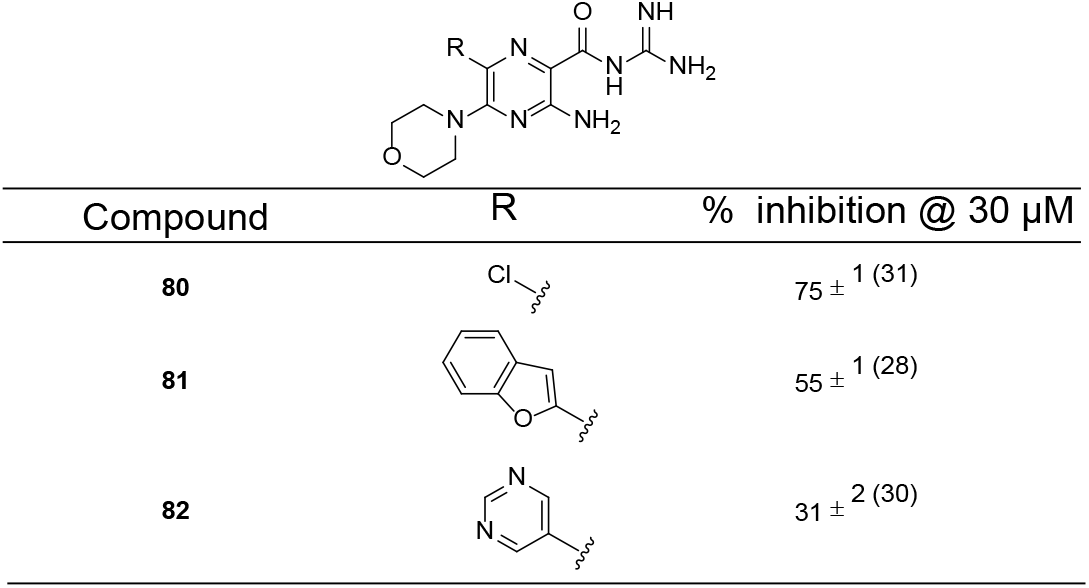
Analogs containing 5-morpholino groups at position 5.

|  |  |  |
| --- | --- | --- |
| Compound | R | % inhibition @ 30 μM |
| **80** | | 75 ± 1 (31) |
| **81** | | 55 ± 1 (28) |
| **82** | | 31 ± 2 (30) |

**Table 8.**
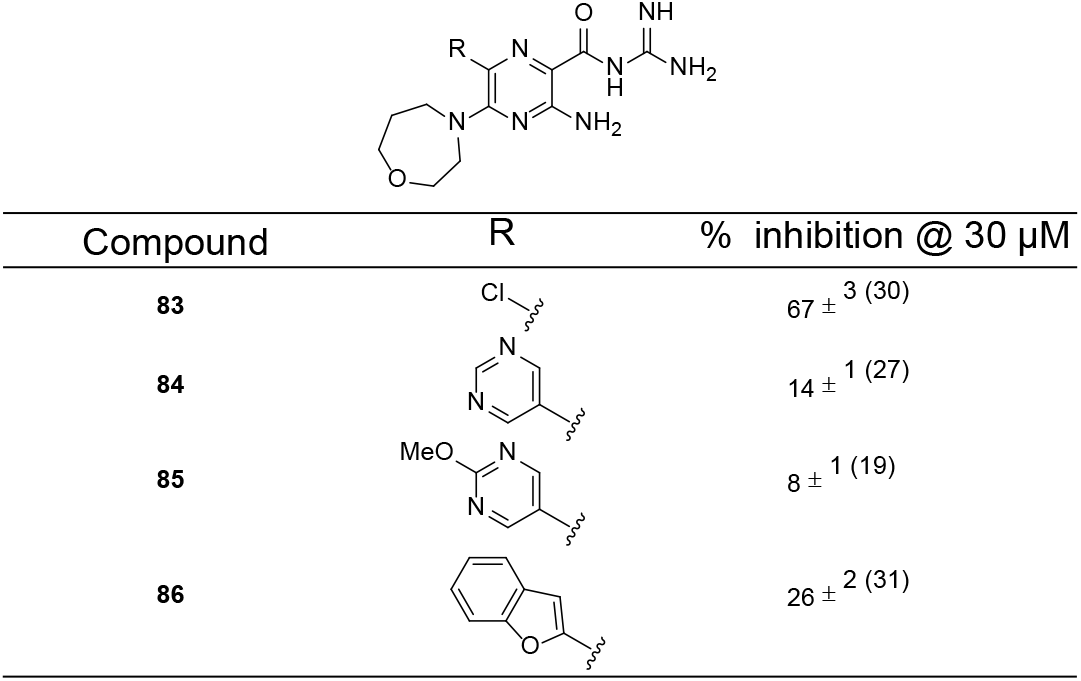
Analogs containing1,4-oxazepane groups at position 5.

|  |  |  |
| --- | --- | --- |
| Compound | R | % inhibition @ 30 μM |
| **83** | | 67 ± 3 (30) |
| **84** | | 14 ± 1 (27) |
| **85** | | 8 ± 1 (19) |
| **86** | | 26 ± 2 (31) |

Screening of 5-*N*-alkyl substituted amiloride analogs (**Table 9**) ^48 48 48 48 48^ confirmed the findings of Kuduk *et al.^37^*, where benzylamine-**87** and phenethylamine-**88** substituted analogs showed higher activity than amiloride (90% and 91% inhibition, respectively). Cyanoethylamino **89**, phosphonic acid ethylamino **90** and *N*-tosyl ethylamino **91** analogs all showed slightly lower activity than amiloride.

**Table 9.**
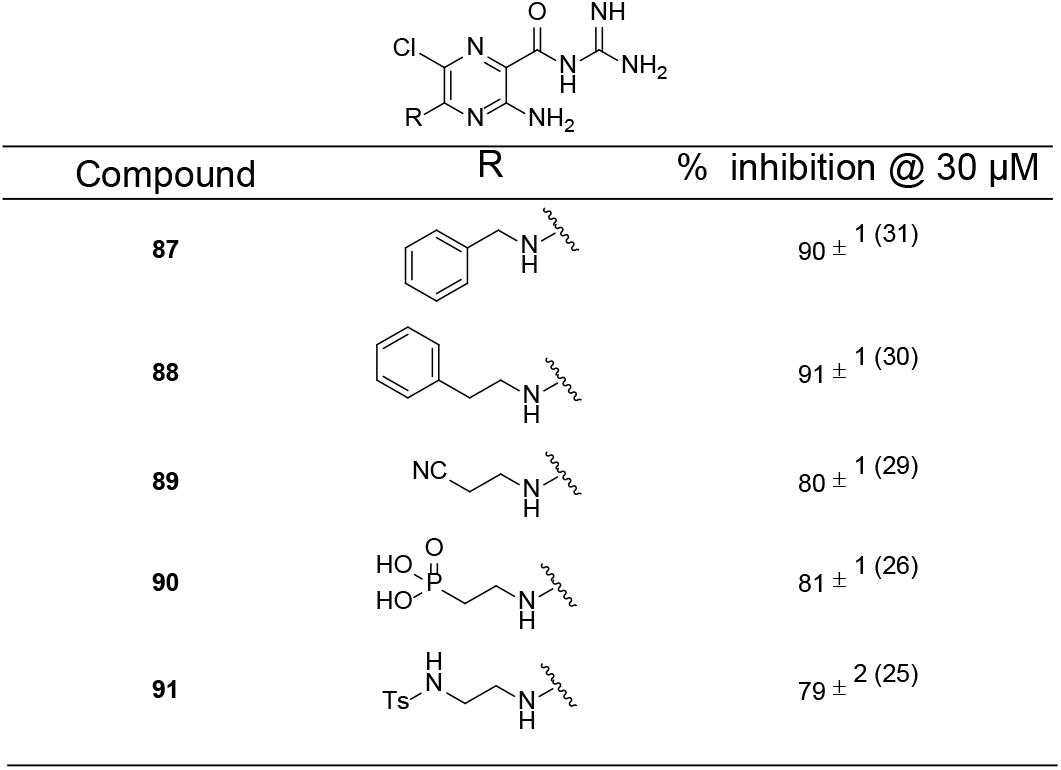
Amiloride analogs containing substituted amines at position 5.

|  |  |  |
| --- | --- | --- |
| Compound | R | % inhibition @ 30 μM |
| **87** | | $90 \pm 1$ (31) |
| **88** | | $91 \pm 1$ (30) |
| **89** | | $80 \pm 1$ (29) |
| **90** | | $81 \pm 1$ (26) |
| **91** | | $79 \pm 2$ (25) |

Simple benzoyl **92**, picolinoyl **93** and nicotinoyl guanidines all showed only low activity at 30 μM (**Table 9**). Slightly higher activity was seen with the aroylguanidine cariporide **94**, a potent and well-studied inhibitor of the sodium-hydrogen exchanger isoform 1-(NHE1).^49^ The guanidine-substituted amiloride analog phenamil **95** was found to be more potent than amiloride **11**, in agreement with a previous finding^30^. With the majority of analogs screened showing similar or reduced activity relative to amiloride, we were delighted to discover that 6-iodo amiloride **97** showed better activity, inhibiting ASIC1 activity by 95% (at 30 μM).

Concentration-response curves were used to determine inhibition constants for a selection of compounds that showed inhibitory activity at 30 μM (**Table 11**). Under our experimental conditions, the estimated *K_i_* (1.7 ± 0.7 μM, n = 5) for amiloride inhibition of ASIC1 currents in tsA-201 cells was in close agreement with and IC_50_ value of 2.2 μM determined by manual patch-clamp (MPC) on HEK-293 cells.^44^ HMA **13** was ~2-fold less potent at inhibiting ASIC1 currents (*K_i_* 4.0 ± 1.9 μM, n = 5), whereas for the 2-benzofuran (**16** and **17**), and 4-methoxypyrimidine 6-substituted amiloride **46** and HMA **47** matched-pairs, the analog containing an unsubstituted amine at position 5 were between 3 and 5-fold more potent than the corresponding 5-azepane compound (**16** *K_i_* = 2.4 ± 0.2 μM, n = 3, cf. **17** *K_i_* = 13.1 ± 2.9 μM, n = 5; **46** *K_i_* = 9.8 ± 5.9 μM cf. **47** *K_i_* = 33.7 ± 23.7 μM). The 5-benzylamine substituted amiloride **87** showed an almost identical *K_i_* to amiloride, according with the single concentration screening data. Addition of a 4-F substituent to the benzyl group **87**, as in **98**, caused a 2.5-fold drop in potency, while further decreases were seen for the 5-(2-cyanoethyl)amino **89** and 5-phenyleythlamino **88** derivatives. *N*-tosyl-substituted ethylenediamine derivative **91** was equipotent with amiloride **11**, while 5-hydroamiloride **99** (**Table S1**) was about 2-fold less active. Interestingly, 6-iodo-amiloride **97** showed a 19-fold increase in potency (*K_i_* = 0.088 ± 0.02 μM, n = 7), revealing it as the most potent amiloride-based ASIC1 inhibitor reported to date.

**Table 10.**
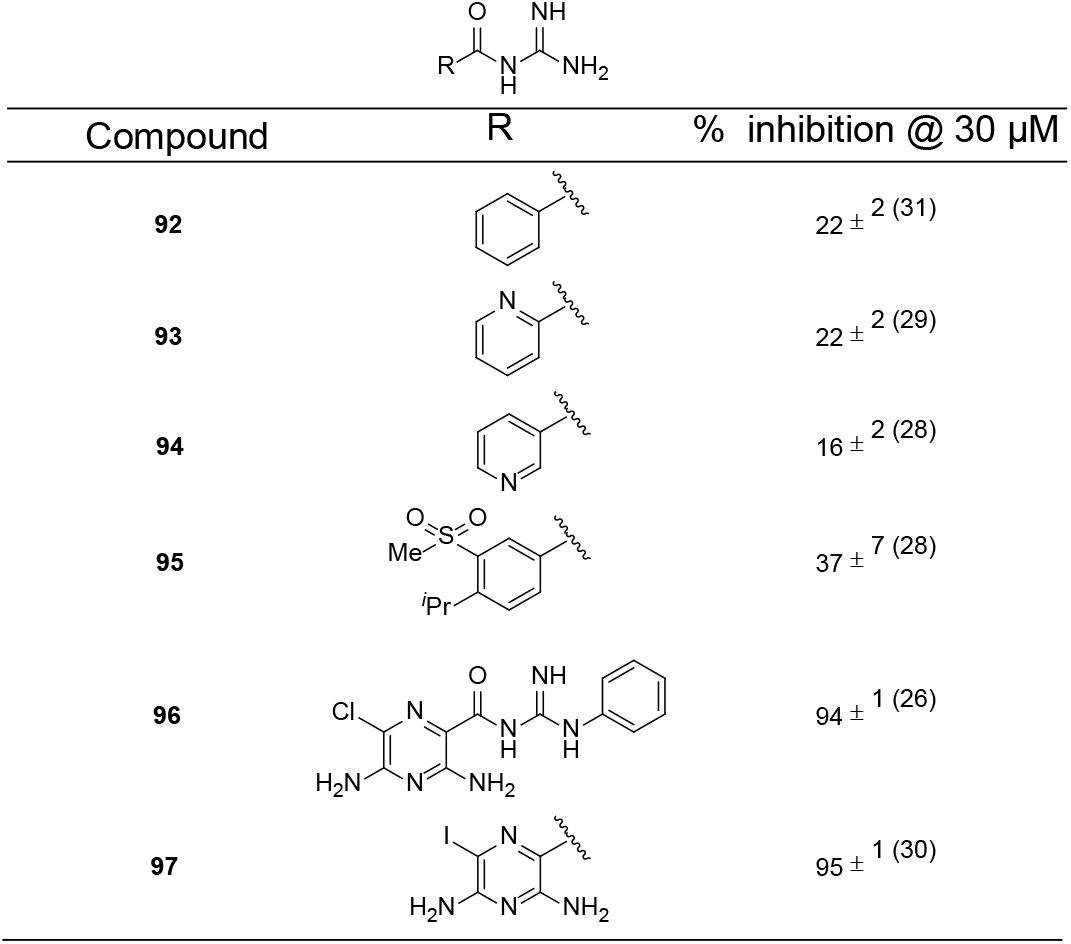
Miscellaneous aroylguanidines and 6-iodoamiloride.

|  |  |  |
| --- | --- | --- |
| Compound | R | % inhibition @ 30 μM |
| **92** | | $22 \pm 2$ (31) |
| **93** | | $22 \pm 2$ (29) |
| **94** | | $16 \pm 2$ (28) |
| **95** | | $37 \pm 7$ (28) |
| **96** | | $94 \pm 1$ (26) |
| **97** | | $95 \pm 1$ (30) |

**Table 11.** ASIC1 inhibition *K_i_* for selected amiloride analogs. Values represent the mean ± SEM with number of technical replicates shown in parentheses. Representative concentration-response curves are show in **Fig S3,4**

| Compound | R<sub>1</sub> | R<sub>2</sub> | IC<sub>50</sub> (μM) |
| --- | --- | --- | --- |
| 11 |  |  | 1.7 ± 0.7 (5) |
| 13 |  |  | 4.0 ± 1.9 (5) |
| 16 |  |  | 2.4 ± 0.2 (3) |
| 17 |  |  | 13.1 ± 2.9 (5) |
| 46 |  |  | 9.8 ± 5.9 (4) |
| 47 |  |  | 33.7 ± 27.3 (5) |
| 87 |  |  | 1.4 ± 0.1 (5) |
| 98 |  |  | 3.5 ± 0.6 (4) |
| 88 |  |  | 4.2 ± 0.6 (3) |
| 89 |  |  | 4.9 ± 0.6 (8) |
| 91 |  |  | 1.8 ± 0.7 (4) |
| 99 |  |  | 3.8 ± 0.8 (4) |
| 97 |  |  | 0.088 ± 0.052 (4) |

To demonstrate the utility of **97** as an ASIC inhibitor in primary tissue we determined its activity in rat dorsal root ganglion (DRG) neurons. DRGs display robust proton-activated currents primarily mediated by ASIC3 expression. While inward currents from DRG neurons elicited by *p*Hs = 6.0 were strongly decreased by 0.3 μM **97** (mauve) and 3 μM amiloride **11** (blue), exposure to 3 μM **97** (pink trace over baseline) completely inhibited all the *p*H 6.0-sensitive current in these cells (**Figure 6A**). Single concentration half inhibitory concentration determined from residual currents during MPC experiments in rat DRG neurons revealed IC_50_ estimates of 0.23 ± 0.02 μM (n = 5) for **97** and 2.72 ± 0.36 μM (n = 5) for **11**, highlighting >10-fold higher potency in neuronal tissue relative to amiloride (**Figure 6B**). Consistent with previous reports,^50^ immunofluorescence confocal micrographs of dorsal root ganglion sections display robust signal from an ASIC3-specific antibody (**Figure 6C**), supporting ASIC3 as a primary mediator of proton-sensitive currents in rat DRG neurons.

**Figure 5.**
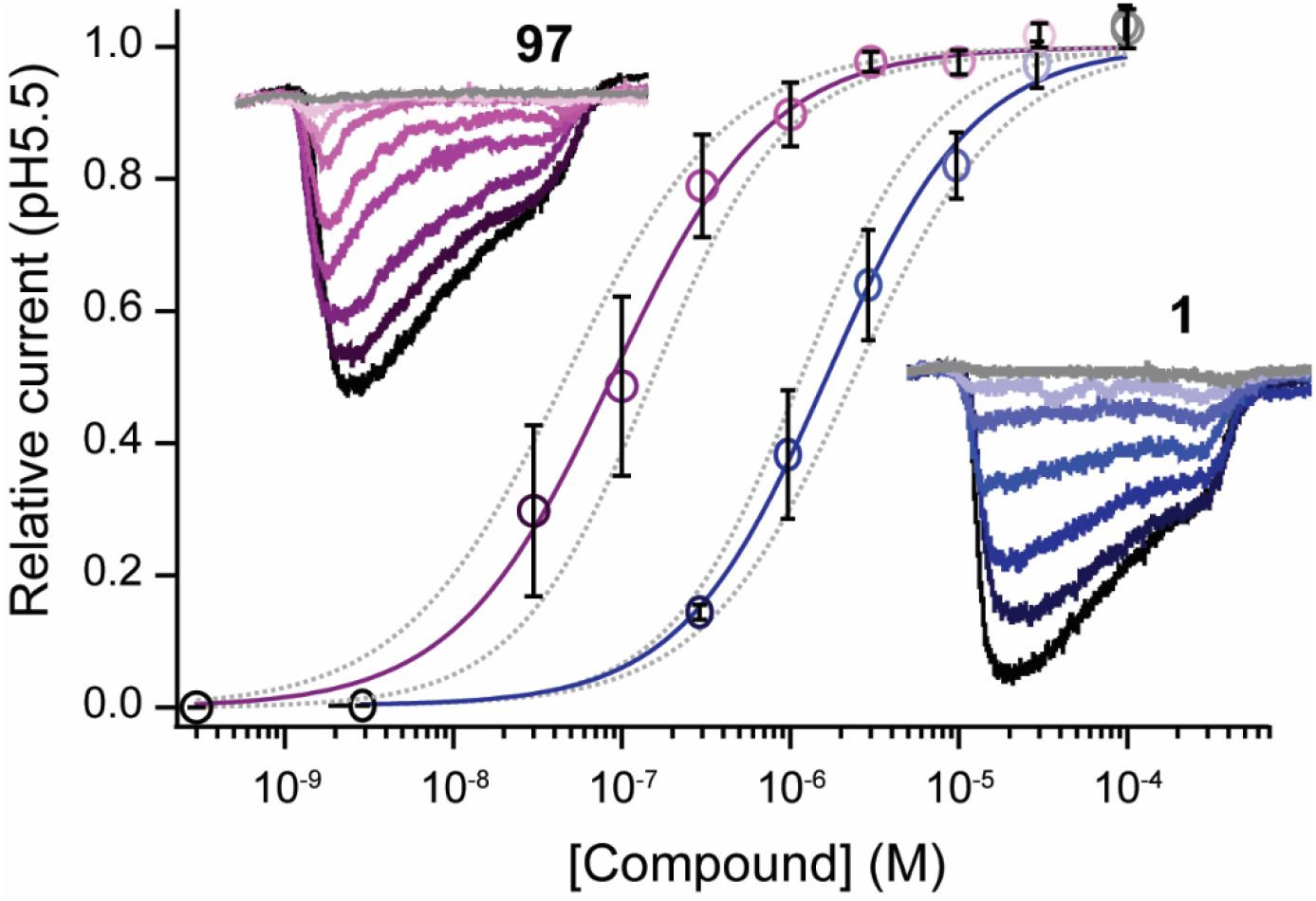
Concentration dependence of ASIC1 current inhibition by **A) 97** (6-iodo-amiloride, mauve; *K_i_* = 0.088 ± 0.02 μM, n = 7) and **B) 11** (amiloride, blue; *K_i_*: = 1.7 ± 0.3 μM, n = 5). The insets show representative currents activated by a *p*H 5.5 jump in the presence of low (dark) to progressively higher (light) [Compound] on tsA-201 cells.

**Figure 6.**
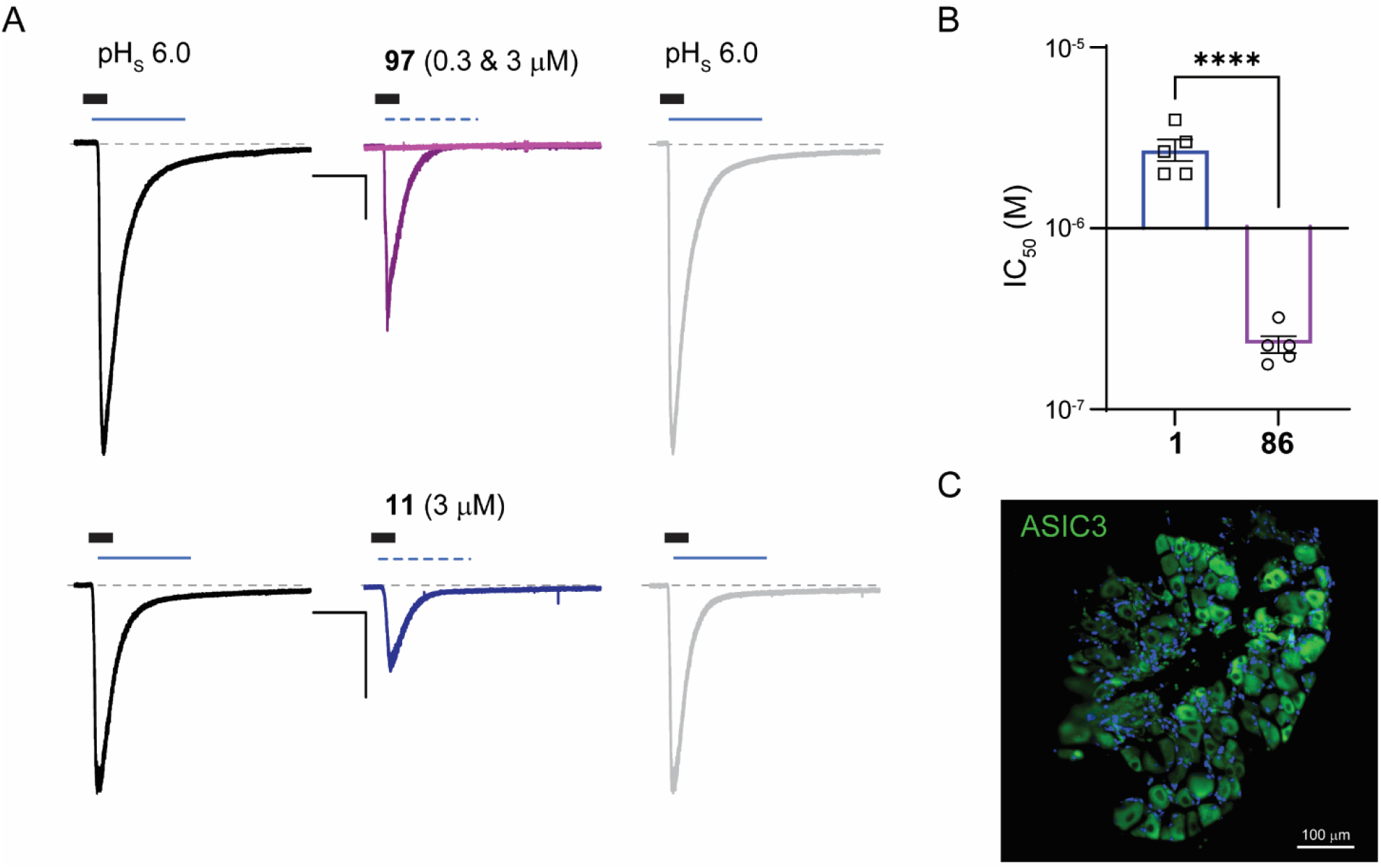
6-Iodoamiloride **97** potently inhibits proton-sensitive currents in rat dorsal root ganglion neurons **A)** Top: inward currents elicited by exposure to extracellular solution at *p*H 6.0 (black) are dose-dependently and reversibly inhibited by **97** (mauve) and amiloride (blue). The exposure time to different treatment was 1 s and is marked by thin lines above the current traces (dotted line marks the application of inhibitor together with the *p*H_s_. Control currents elicited by *p*H 6.0 and after inhibitor washout are shown in black and light grey, respectively. Thick black bars mark the conditioning *p*H_C_ = 7.4. Scale bars are 10 s and 0.5 nA. **B)** Half inhibitory concentrations calculated from residual currents DRG neuron proton-gated currents in the presence of **97** (0.3 μM) and amiloride **11** (3 μM). Single concentration IC_50_s **97** = 0.23 ± 0.02 μM (n = 5) and **11** = 2.72 ± 0.36 μM (n = 5), unpaired *t*-test p = 0.0001. **C)** Immunofluorescence confocal micrograph of a section of dorsal root ganglion showing robust signal from an ASIC3-specific antibody (green) and counterstained with DAPI (blue nuclei).

### Conclusions

This work describes novel SAR for the amiloride class against ASIC-mediated H^+^-gated currents in whole cells and primary neuronal tissues, providing insight into its suitability as starting point for the medicinal chemistry optimisation of new ASIC inhibitors. Relatively flat SAR was seen at the 6-position of the amiloride pyrazine core for a diverse range of (het)aryl analogs with increases in potency not able to be achieved. Activity was further decreased by additional (cyclo)alkyl substitution at the 5-position for sub-series of 6-substituted analogs, indicating a lack of compatibility for substitutions at these positions. Trends favouring the introduction of larger secondary alkylamines at the 5-position within the amiloride (6-Cl) series were consistent with previous reports for other amiloride analogs at this position. To our surprise, screening of the previously described amiloride analog, 6-iodoamiloride **97** revealed a robust improvement in potency, with a *K_i_* in the sub-100 nM range. 6-iodoamiloride represents the most potent amiloride-derived ASIC inhibitor reported to date, while also being the most potent competitive small molecule ASIC inhibitor yet described. Disease-relevant inhibition of H^+^-activated currents was demonstrated in rat primary DRG neurons. 6-iodoamiloride may prove a useful tool for further dissecting the roles of ASIC function in various disease models.

## Methods

### Cell culture

tsA-201 cells were grown at 37 °C and 5% CO_2_ in DMEM supplemented with 10% FBS (Bovogen, France), 2 mM GlutaMAX, 100 U/mL penicillin, 10 μg/mL streptomycin. Cells were enzymatically dislodged and passaged every 2-3 days. On the day of the experiment, cells monolayers were detached using TrypLE (ThermoFisher Scientific) and re-suspended at 100,000 cells/mL in cold ChipFill solution. Cells were maintained in suspension Cell Hotel reservoir at 12 °C with shaking at 200 rpm.

### Recording solutions and compound preparation

For recordings, the standard intracellular solution (IS, 60 mM CsF, 50 mM CsCl, 10 mM NaCl, 20 mM EGTA and 10 mM HEPES (*p*H 7.2, adjusted with CsOH, 280 ± 3 mOsm); and extracellular solution (ES,140 mM NaCl, 4 mM KCl, 2 mM CaCl_2_, 1 mM MgCl_2_, 5 mM Glucose and 10 mM HEPES; *p*H 7.4, adjusted with NaOH, 298 ± 3 mOsm). The *p*H of the ligand solution (ES at *p*H 5.5) was buffered with 10 mM 2-(*N*-morpholino)ethanesulfonic acid. The chip fill solution 140 mM NaCl, 4 mM KCl, 3 mM MgCl_2_, 5 mM glucose, and 10 mM HEPES (*p*H 7.4, adjusted with NaOH, 290 ± 3 mOsm). Seals were formed by brief treatment with seal enhancement solution (SE, 80 mM NaCl, 3 mM KCl, 10 mM MgCl_2_, 35 mM CaCl_2_ and 10 mM HEPES, *p*H 7.3, adjusted with NaOH, 298 ± 3 mOsm) and washed out thoroughly with standard external solution. All solutions were 0.22-μm membrane filtered and compounds were dissolved in DMSO as 100 mM stock solutions and diluted appropriately to yield the final test concentrations. Final DMSO concentrations were maintained below 0.1% v/v.

### Automatic Planar Patch-Clamp Electrophysiology

Single-concentration compound screening was performed using a SyncroPatch 384PE module (Nanion Technologies GmbH, Germany) incorporated into a Biomek FX pipetting robot (Beckman Coulter, Jersey City, NJ). Recordings were carried out in planar borosilicate glass patch clamp chips in a 384-well microtiter plate format with medium-resistance, eight-hole-per-well chips # 221801 (Nanion Technologies GmbH). Concentration-response experiments were performed using an NPC-16 Patchliner (Nanion Technologies, Munich, Germany) using multi-hole medium resistance NPC-16 chips # 071401 with an average resistance of 1.1 MΩ. Whole-cell patch-clamp voltage-clamp experiments were performed at room temperature (21-23 °C). Chip and whole-cell capacitance were fully compensated, and series resistance compensation (70%) was applied via the “Auto Rs Comp function”. Currents were evoked by ~1 s puff application of ligand (ES *p*H 5.5 or *p*H 5.5 + 30 μM compounds) during a 5 s test pulse from a holding potential of −70 mV. Compound was applied to each cell and exposed for 240 s before ligand + compound test puff.

### Data Analysis

Data acquisition and analysis was performed with PatchControl 384 and DataControl 384 software, respectively (Nanion Technologies, Germany). Only cells with a seal resistance R_seal_ > 500 MΩ were included for analysis. A test pulse in control bath solution application (ES, *p*H 7.4) was used to subtract leak currents. Offline analysis was performed using DataControl 384 and Igor Pro-6.37 software (WaveMetrics Inc., Portland, OR, USA). The integer of area under the curve described by the inward currents in response to the ligand puff was used for quantification of agonist-induced responses.

### Dorsal root ganglion isolation and culture

Animal studies are reported in compliance with the ARRIVE guidelines (Kilkenny, Browne, Cuthill, Emerson, & Altman, 2010). DRGs were isolated from neonatal (P12-P14) Sprague Dawley rats (ArcCrl:CD (SD)IGS, Animal Resources Centre, Australia, RRID:RGD_8553001) following decapitation, as approved by the University of Wollongong Animal Ethics Committee (AE17/23). Rats were housed in double decker, individually ventilated cages containing corncob bedding, nesting material, chewing block, PVC tunnel, and a plastic or cardboard shelter. Rats were housed at 21 °C on a 12 h light/dark cycle, and fed Vella Rat Pellets and water *ad libitum*. Briefly, DRGs were collected by hemisecting the spinal cord and removing individual ganglia, in ice-cold divalent-free Hank’s balanced salt solution (HBSS) where dorsal root and peripheral nerve processes were carefully trimmed using spring scissors (Fine Science Tools, Canada). Ganglia were transferred to dissociation media (divalent cation-free HBSS with 2 mg/mL collagenase Type II (CLS2, Worthington, USA) for 60 min at 37 °C, 5% CO_2_. Digested ganglia were centrifuged at 200 g and 23 °C for 10 min and the supernatant discarded. The isolated neurons were washed with DMEM (supplemented with 10% FBS, 1× GlutaMAX, and 1% penicillin/streptomycin) followed by trituration with progressively smaller diameter fire polished Pasteur pipettes. Dissociated neurons were then filtered through a 160 μm nylon mesh (Millipore, Australia) to remove cell debris and non-dissociated material. The neuron suspension was then plated on poly-*L*-lysine and laminin coated 12 mm cover glass (Sigma, Australia) and left to attach for ~3 h at 37 °C, 5% CO_2_ after which 1 mL of fresh DMEM (10% FBS, 1× GlutaMAX, and 1% pen/strep) was added and incubated overnight. Primary DRG cultures were kept at 30 °C and recorded within 48 h of isolation.

### Manual patch clamp

Whole-cell patch clamp recording of rat dorsal root ganglion neurons were carried out 1–2 days post-isolation. Neurons were perfused with extracellular solution containing 140 TEA-Cl, 10 CaCl_2_, 1 MgCl_2_, 10 D-glucose, 10 HEPES, and *p*H 7.4 with TEA-OH (~310 mOsmol/kg). Borosilicate patch pipettes (World Precision Instruments, USA) were fire-polished to a resistance of 1–3 MΩ and filled with intracellular solution containing (mM): 150 CsCl, 1.5 MgCl_2_, 5 EGTA, 10 HEPES, and *p*H 7.2 with CsOH (~300 mOsmol/kg). H^+^-activated currents (*p*H 6.0) were recorded at room temperature (22–24°C) using a MultiClamp 700B amplifier (Molecular Devices, USA), digitalized via a Digidata 1440 controlled by pClamp10.7 acquisition system. Compounds were superfused to the neuron using a syringe pump (New Era Pump Systems Inc, USA) loaded with 1 mL syringe connected to a 50 cm MicroFil 28G (World Precision Instruments) at 2 μL/min allowing rapid solution exchanges.

### Immunostaining and confocal imaging

tsA-201 cells at ~50% confluence and dissociated DRG neurons were fixed with 4% w/v paraformaldehyde/PBS without Mg^2+^ or Ca^2+^ (PBS^Div-^) for 15 min, washed 3 times with PBS counterstained with wheat germ agglutinins 0.01 mg/mL for 30 min at RT. Whole DRGs were fixed overnight in 4% paraformaldehyde/PBS^Div-^(*p*H 7.3 with NaOH) and cryoprotected in 30% w/v sucrose/PBS^Div-^for 48 h. Ten sagittal sections of 20 μm thickness were cut through the entire DRG using a cryostat (Leica, Richmond, IL, USA) and placed on poly-*L*-lysinated glass slides, air-dried overnight and rehydrated with ddH_2_O. All samples were then permeabilized with 0.1% v/v Triton X-100/PBS^Div-^for 15 min and blocked in 1% v/v normal goat serum/0.1% v/v Triton X-100/PBS^Div-^for 30 min at RT. All primary antibodies were diluted at the concentrations indicated in 1% v/v normal goat serum/0.1% v/v Triton X-100/PBS^Div-^.Antibody-free diluent was used as negative control. Alpha Diagnostic International (San Antonio, TX, USA) ASIC Pan 1-3 Mab cocktail (1:500; ASIC-PAN-51-A) and ASIC1a rabbit anti-rat (1:100; ASIC11-A) primary antibodies were incubated for 1 h at RT, followed by two washes in 1% v/v Triton X-100/PBS^Div-^. Cells were then exposed to Alexa-Fluor 488 or 568-tagged secondary goat anti-rabbit Ig (1:200, Molecular Probes, Eugene, OR, USA) for 30 min at RT. Slides were then rinsed with ddH_2_O, and coverslips were applied to tissue sections with ProLong Gold antifade reagent with 4,6”-diamidino-2-phenylindole (DAPI; Invitrogen) to stain nuclei and protect fluorescence from fading. Nuclei were then stained with 0.01% w/v 4’,6-diamidino-2-phenylindole dilactate (DAPI, Invitrogen) for 5 min and rinsed with ddH_2_O, and coverslips were mounted with Fluorescence Mounting Medium. All images were acquired with Leica TCS SP8 confocal microscope with Tandem Scanner 12 kHz, VISIR scan optic with rotation, two Internal Detector Channels (PMT) and one HyD source, equipped with four lasers: laser 405 nm DMOD Compact; blue 488 nm, green 552 nm, red. All images were obtained in sequential scanning laser mode to avoid fluorochrome cross-excitation (A Z-stack of 15–30 optical). Appendix A. Supplementary data

## AUTHOR INFORMATION

B.J.B. conceived the study, reviewed all data and wrote the manuscript. R.K.F.-U. designed, performed and analyzed all APC experiments and co-wrote the manuscript. B.J.B., A.A., R.S.B. and H.M. synthesized and characterized the compounds. J.R.M. performed electrophysiology experiments with primary rat DRG neurons. A.R. performed the immunostaining experiments. D.J.A., M.K. and M.R provided advice and reviewed the manuscript. All authors approved the final manuscript.

## Funding Sources

Funded in part by an Illawarra Health and Medical Research Institute Young Investigator Grant to B.J.B., Rebecca Cooper Foundation for Medical Research Project Grant (PG2019396) to J.R.M., and Australian National Health and Medical Research Council (NHMRC) Project Grant (APP1100432) to M. J. K. and M.R.

## ACKNOWLEDGMENTS

B.J.B. Gratefully acknowledges salary support from the Illawarra Cancer Carers.

## ABBREVIATIONS

## Supplementary Information

**Figure S1.**
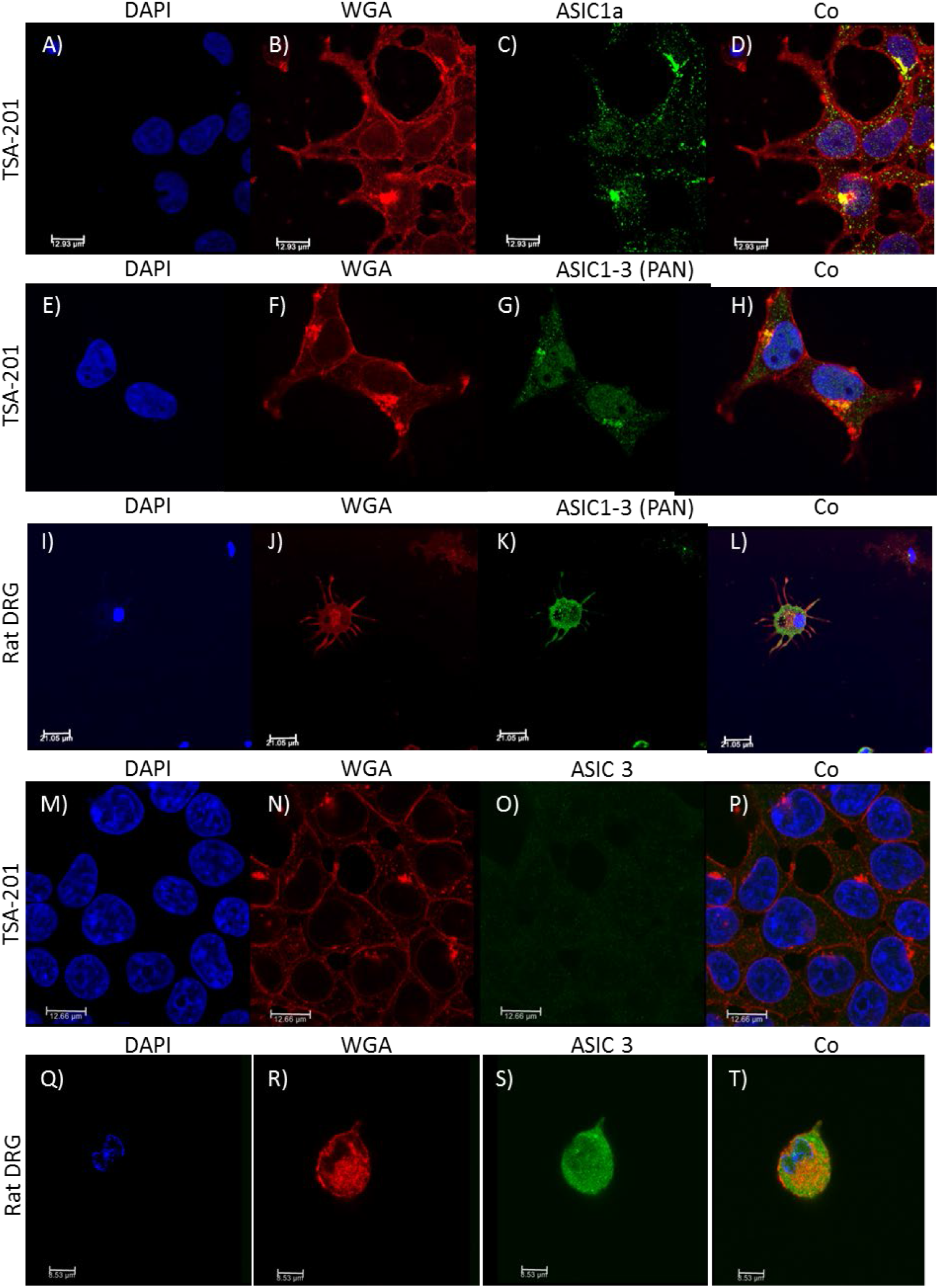
Immunofluorescence images of tsA-201 cells and rat dorsal root ganglion (DRG) neurons stained for ASIC1a and ASIC3 channels (counterstaining: membrane WGA-594; nuclei DAPI). Confocal images for channel are provided. Bottom: the rightmost panel displays an overlay of ASIC3 (green) and WGA-594 (red).

**Figure S2.**
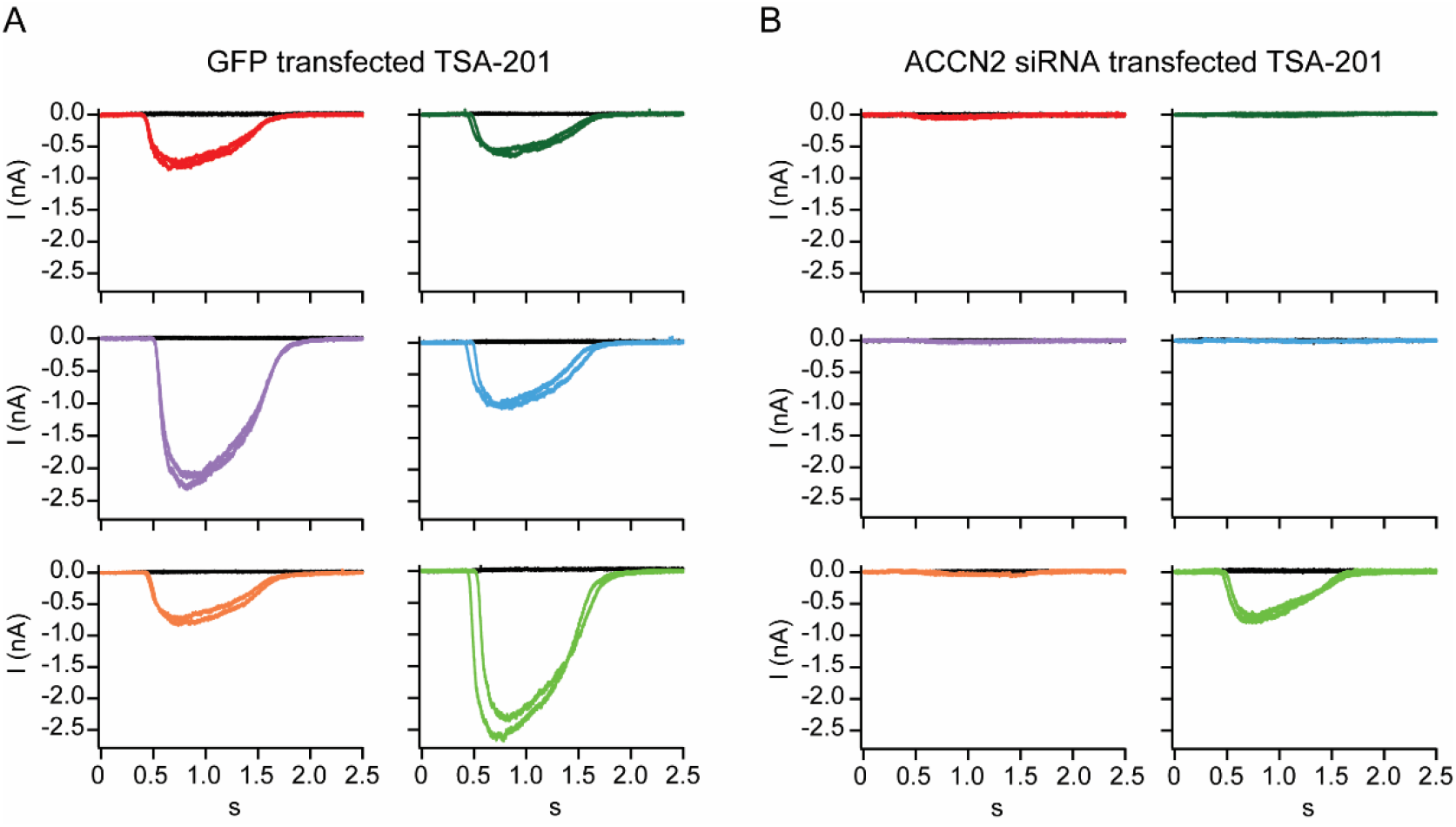
Knockdown of ASIC1 encoding transcripts in tsA-201 cells eliminates proton-gated currents in transfected cells. Current were elicited by *p*H 5.5 exposure of tsA-201 cells transfected with GFP alone **A)** or in combination with anti-ACCN2 siRNAs **B)**. A transfection efficiency of 83% was estimated from the percentage of GFP+ cells in both groups.

**Figure S3.**
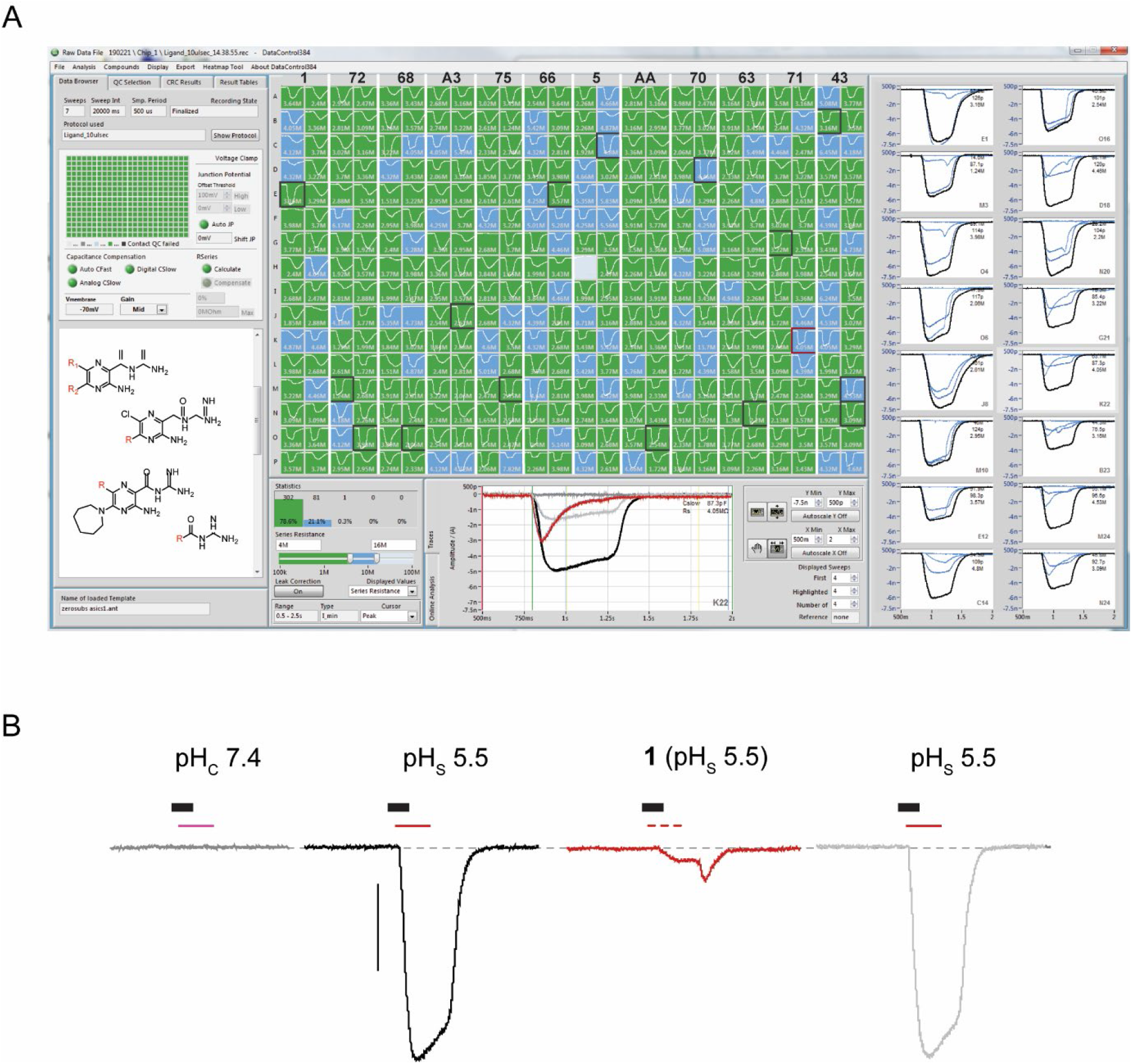
Automated patch clamp screening of library of 6-substituted and 5,6 disubstituted amiloride analogs. **A)** Example experimental setup testing 11 analogs along with reference amiloride **1** (2 columns per analog) and superimposed ASIC1 current traces from tsA-201 cells recorded in the SyncroPatch 384PE and elicited by exposure to pH 7.4, pH 5.5, pH 5.5+analog, and pH 5.5 after washout. **B)** Current traces are shown according to their sequential treatments (scale bar is 1 nA). The conditional extracellular solution (pHC 7.4) is represented by the thick black bar whereas the different treatments are marked by thin lines. In **B** and the bottom of **A** (expanded well K22/**71**, for reference), current responses to puffs of pH 7.4, pH 5.5, pH5.5 + **analog** and pH 5.5 (after compound washout) are shown in dark grey, black, red and light grey, respectively.

**Figure S4.**
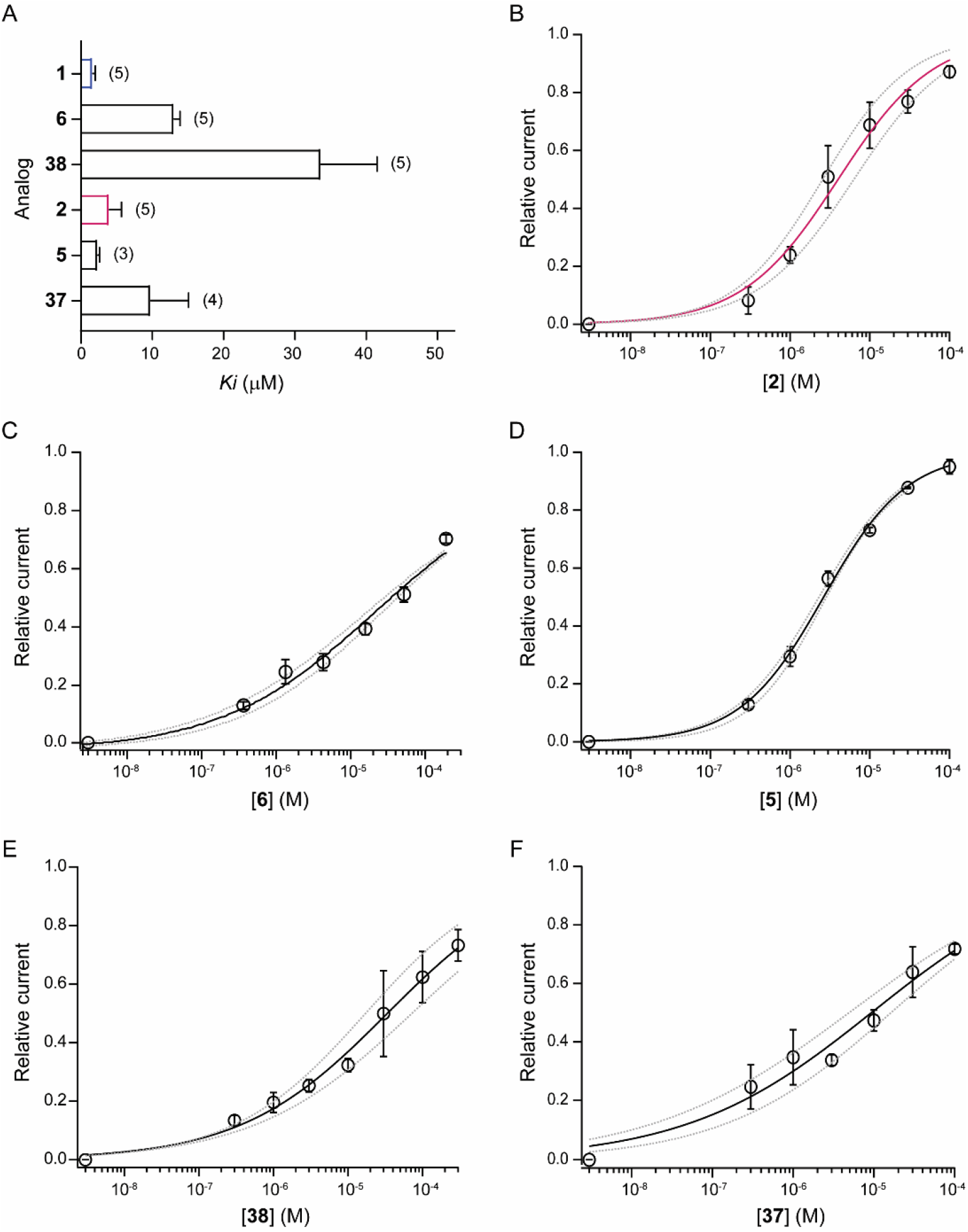
**A)** Summary of affinity determination for amiloride analogues. Concentration-response curves for inhibition of endogenous ASIC-mediated currents in tsA-201 for **B) 12** (*K_i_* = 4.0 ± 1.7 μM, nH =0.73, n = 5); **C) 17** (*K_i_* = 13.1 ± 0.8 μM, nH = 0.45, n = 5); **D) 16** (*K_i_* =2.4 ± 0.2 μM, nH = 0.88, n= 3); **E) 47** (*K_i_* = 33.7 ± 7.8 μM, nH = 0.44, n = 5); and **F) 46** (*K_i_* = 9.8 ± 5.3 μM, nH = 0.37, n = 4).

**Table S1.**
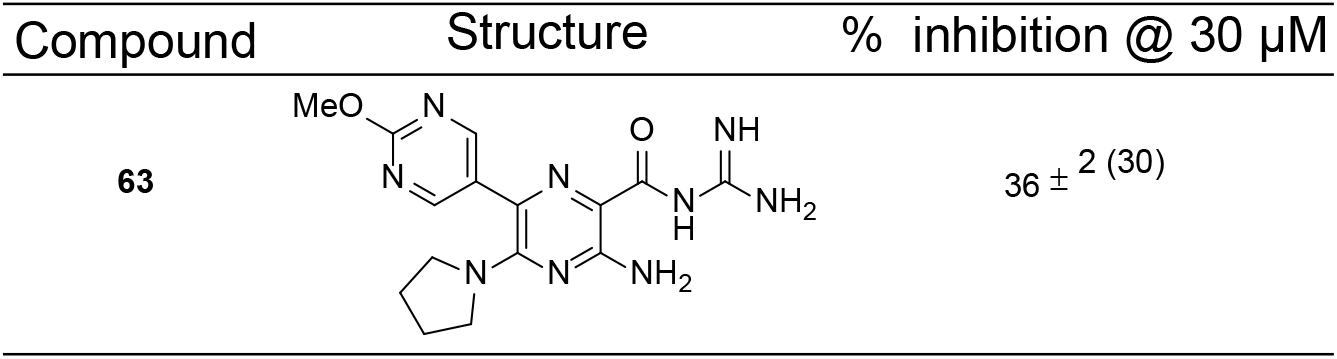

| Compound | Structure | % inhibition @ 30 μM |
| --- | --- | --- |
| **63** | | $36 \pm 2$ (30) |

## Notes

### Competing Interest Statement

The authors have declared no competing interest.

